# The genetic basis of the human-cannabis relationship

**DOI:** 10.1101/287938

**Authors:** Philippe Henry

## Abstract

Cannabis can elicit various reactions in different consumers. In order to shed light on the mechanisms underlying the human-cannabis relationship, we begin to investigate the genetic basis of this differential response. The web-based platform OpenSNP was used to collect selfreported genetic and phenotypic data. Participants either reported a positively or negative affinity to cannabis. A total of 26 individuals were retained, 10 of which indicated several negative responses and the remaining 16 indicating strong affinity for Cannabis. A total of 325’895 single nucleotide polymorphisms (SNPs) were retained. The software TASSEL 5 was used to run a genome-wide association study (GWAS), with a generalized liner model (GLM) and1000 permutations. The analysis yielded a set of 45 SNPs that were significantly associated with the reported affinity to cannabis, including one strong outlier found in the MYO16 gene. A diagnostic process is proposed by which individuals can be assessed for their affinity to cannabis. We believe this type of tool may be helpful in alleviating some of the stigma associated with cannabis use in individuals sensitive to THC and other cannabis constituents such as myrcene, which may potentiate negative responses.

## Background

Cannabis (*Cannabis sativa* L.) is often cited as the most commonly used recreational drug worldwide. The most recent National Survey on Drug Use and Health confirms this assertion, showing that about 34% to 42% of Americans have used cannabis in their lifetime (*one-time users*), 10% to 12% have used it in the past year (*casual users*), 6% to 7% are *regular users* and about 4% are *chronic users* (Substance Abuse and Mental Health Services Administration, 2014). It is interesting to note that these statistics are mirrored in other jurisdictions around the world (e.g. Canada), leading us to hypothesize that a differential affinity (DA) towards cannabis may exist in human populations.

While most participants may experience a feeling of relaxation, exuberance, laughter and/or hunger, others may react differently and exhibit social anxiety or other negative responses. Such outcomes may be mediated by individual differences in gene encoding the endocannabinoid system and other associated regulatory enzymes, along with unexplored regions of the genome.

## Methods

In the present research, an open access data set consisting of nearly 350’000 genome-wide single nucleotide polymorphisms (SNPs) typed in volunteer participants using the web-based platform OpenSNP: (https://opensnp.org/phenotypes/401).

Participants self-reported either a positive or negative affinity to cannabis. After initial screening, a total of 26 out of 53 individual respondents were retained, 10 of which indicated several negative responses and the remaining 16 indicating strong affinity for cannabis. In addition to the genome-wide dataset, two targeted datasets were built. 1) hmnVSSL-a set of 15 SNPs associated with the endocannabinoid system, and 2) 420andme - consisting of SNPs selected using the approach described in the GWAS below.

The software TASSEL 5^1^ was used to run a genome-wide association study (GWAS), with a generalized liner model (GLM) and1000 permutations. TASSEL was also used to generate a principal component analysis (PCA) and a neighbor-joining tree that were visualized using the software R^2^.

## Results and discussion

The analysis yielded a set of 45 SNPs that were significantly associated with the reported affinity to cannabis, including one strong outlier found in the MYO16 gene.

The three datasets were compared in their resulting neighbor-joining trees and PCA outputs as shown in Fig.1. The 45 SNPs from the 420andme dataset outperformed both the genome-wide and the targeted hmnVSSL datasets in term of its ability to discriminate the cannabis affinity phenotype using genetic data.

**Figure 1.**
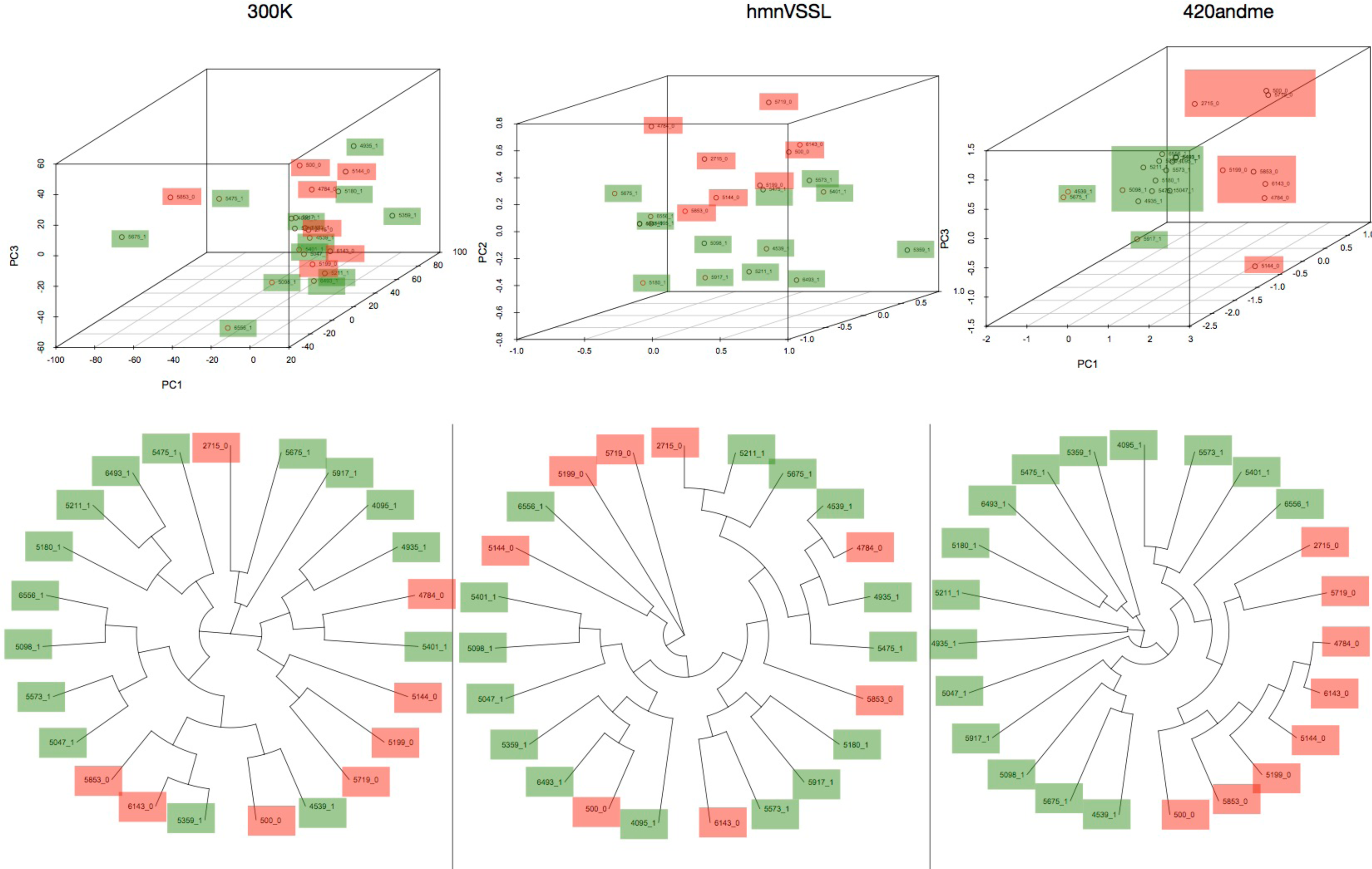
3D Principal Componenet Analysis and Neighbor-Joining trees shown for each of the three datasets: - genome-wide 300 thousand SNPs, - hmnVSSL - 15 SNPs in the endocannabinoid system, - 420andme - 45 SNPs associated with affinity to cannabis phenotypes. One can appreciate the performance of the 420andme dataset in teasing apart human-cannabis affinity phenotypes. Green indicates an individual with a positive affinity for cannabis, whereas red indicates a negative response to cannabis use.

We propose that this type of assays could be run on a qPCR system to discriminate genotypes at the 420andme SNPs in order to predict one’s affinity to cannabis and possibly predict and avoid negative responses. These targeted assays will offer a cost effective manner of typing a large number of participants and could be ideal for clinical and recreational genomics alike.

## Acknowledgements

No formal funding was provided for this study and all analyses were undertaken pro-bono with the intent to develop novel resources for an under-studies system. The individuals who volunteered their genetic and phenotypic data to the project are thanked for their participation and open-ness to this endeavor.

## References

1- Bradbury PJ, Zhang Z, Kroon DE, Casstevens TM, Ramdoss Y, Buckler ES. (2007) TASSEL: Software for association mapping of complex traits in diverse samples. Bioinformatics 23:2633–2635.

2- R Core Team (2018). R: A language and environment for statistical computing. R Foundation for Statistical Computing, Vienna, Austria. URL https://www.R-project.org/.

